# Time-resolved compound repositioning predictions on a text-mined knowledge network

**DOI:** 10.1101/625459

**Authors:** Michael Mayers, Tong Shu Li, Núria Queralt-Rosinach, Andrew I Su

## Abstract

**Background:** Computational compound repositioning has the potential for identifying new uses for existing drugs, and new algorithms and data source aggregation strategies provide ever-improving results via *in silico* metrics. However, even with these advances, the number of compounds successfully repositioned via computational screening remains low. New strategies for algorithm evaluation that more accurately reflect the repositioning potential of a compound could provide a better target for future optimizations.

**Results:** Using a text-mined database, we applied a previously described network-based computational repositioning algorithm, yielding strong results via cross-validation, averaging 0.95 AUROC on test-set indications. The text-mined data was then used to build networks corresponding to different time-points in biomedical knowledge. Training the algorithm on contemporary indications and testing on future showed a marked reduction in performance, peaking in performance metrics with the 1985 network at an AUROC of .797. Examining performance reductions due to removal of specific types of relationships highlighted the importance of drug-drug and disease-disease similarity metrics. Using data from future timepoints, we demonstrate that further acquisition of these kinds of data may help improve computational results.

**Conclusions:** Evaluating a repositioning algorithm using indications unknown to input network better tunes its ability to find emerging drug indications, rather than finding those which have been withheld. Focusing efforts on improving algorithmic performance in a time-resolved paradigm may further improve computational repositioning predictions.

## Background

Compound repositioning is the identification and development of new uses for previously existing drugs. Repositioning is an attractive pipeline for drug development primarily due to the reduced pharmaceutical uncertainty and development times when compared to traditional pipelines [1]. While clinical observation and improved understanding of the mechanism of action are the two primary means by which a drug is repositioned, computational repositioning provides a third route to identifying these candidates. This third method has seen much development in the past decade as a way to potentially speed up the drug discovery process. The ultimate goal of computational repositioning is to quickly produce a small number of clinically relevant hits for further investigation. This process is achieved through the identification of features that relate drugs to diseases and utilizes a gold standard of known true drug-treats-disease relationships to train an algorithm to categorize or rank potential drug-disease pairs for treatment probability. While this path can efficiently produce repositioning probabilities for countless drug-disease pairs, identifying and experimentally validating the results of clinical importance can be both costly and challenging [2].

In the last decade, there have been many improvements in approaches and algorithms to identify these candidates [3]. These include an expansion from gene expression-based approaches [4, 5] to include methods based on knowledge graphs [6, 7]. Coupled with the advancements in machine learning, the number of different methods for producing repurposing predictions has quickly increased, each showing marked improvements on their ability to accurately predict candidates. One common result in these knowledge-based approaches is that drug-drug and disease-disease similarity, when combined with drug-disease associations, provide the important information for generating a learning model [6, 8, 9]. Many different metrics can be used to express these similarities, like structural motifs in the case of drugs, or phenotypes in the case of diseases. However, as good as these algorithms have become at providing repurposing candidates from a list of known indications, the majority of computational repositioning projects do not continue beyond the *in vitro* studies [10].

One recent effort in computational repositioning, Himmelstein et. al.’s Rephetio project [11] used a heterogeneous network (hetnet) to describe drug-disease relationships in a variety of ways. This method worked by extracting counts of various metapaths between drug-disease pairs, where a metapath is defined by the concept and relationship types in the knowledge graph that join the drug and disease. These metapaths counts are then used as numerical features in a machine learning model. This study compiled several different highly curated data sources to generate the hetnet underlying this learning model and achieved excellent performance results. Whether this learning model that utilizes network structure as features can achieve similar results with a less well-curated network remains an open question.

Progress in the field of natural language processing (NLP) has led to the ability to generate large biomedical knowledge bases through computational text-mining [12, 13]. This method can produce large amounts of data rather quickly, which when coupled with semantic typing of concepts and relations, produces a massive datasource that can quickly be represented in a hetnet structure.

In this work, we evaluated the utility of text-mined networks for use in computational compound repositioning, by utilizing the Semantic MEDLINE Database (SemMedDB) [14] as an NLP-derived knowledge network, and the Rephetio algorithm for producing predictions. We evaluated the performance of this data source when trained with a gold standard of indications taken from DrugCentral [15] and tested via cross-validation. We then propose a new framework for evaluating repurposing algorithms in a time-dependent manner. By utilizing one of the unique features of SemMedDB, a PubMed Identification number (PMID) documented for every edge in the network, multiple networks were produced in a time-resolved fashion, each with data originating on or before a certain date, representing the current state of knowledge at that date. These networks were then evaluated in the context of computational repositioning via training on indications known during the time period of the given network and tested on indications approved after the network, a paradigm that more closely resembles the real-world problem addressed by computational repositioning than a cross-validation. Finally, we analyzed these results to identify the types of data most important to producing accurate predictions and tested the predictive utility of supplementing a past network with future knowledge of these important types.

## Methods

### Initial SemMedDB Network Generation

The SemMedDB SQL dump Version 31R, processed through June 30, 2018, was downloaded (https://skr3.nlm.nih.gov/SemMedDB/download/download.html) and converted into a csv. Using Python scripts (https://github.com/mmayers12/semmed/tree/master/prepare), corrupted lines were removed, and lines were normalized to a single subject-predicate-object triple per line, with identifiers in Unified Medical Language System (UMLS) space. This ‘clean’ database was then further processed into a heterogeneous network (hetnet) compatible with the hetnet package, hetio (https://github.com/hetio/hetio) a prerequisite for the rephetio machine learning pipeline. This processing included: using the UMLS Metathesaurus version 2018AA to map terms to other identifier spaces (primarily Medical Subject Headings or MeSH), combining granular concepts into a more general terms, thus reducing node-count and data-redundancy; combining semantic (edge) types of similar meaning (e.g. between Chemicals & Drugs and Disorders, ‘TREATS’, ‘PREVENTS’, ‘DISRUPTS’, and ‘INHIBITS’ were merged to ‘TREATS’); filtering out semantic edge types that were sparsely populated (less than 0.1% of the total network); removing the top 100 nodes by degree to eliminate extremely general concepts (e.g., Patients, Cells, Disease, Humans); filtering out edges with less than 2 supporting PMIDs to reduce data noise due to text-mining.

To create time-resolved knowledge networks, a map between PMID and publication year was generated from four data sources: Pubmed Central (ftp://ftp.ncbi.nlm.nih.gov/pub/pmc/), Euro PMC (http://europepmc.org/ftp/pmclitemetadata/), NLM - Baseline Repository (ftp://ftp.ncbi.nlm.nih.gov/pubmed/baseline/), and EBI’s API (https://europepmc.org/RestfulWebService). Output from these sources was merged to encompass the greatest number of PMIDs possible. Networks were generated at 5-year intervals starting at the year 1950 continuing to present day. The PMID with the earliest publication year for a given edge was used for that edge.

### Gold standard generation

The PostgreSQL dump of DrugCentral dated 2018-06-21 was downloaded for use as the gold standard of known drug-disease indications. The following tables were extracted for use throughout the analysis pipeline: *omap_relationship*, containing the indications; *identifier*, with maps from internal IDs to other systems including UMLS and MeSH; *approval*, containing approval dates from worldwide medical agencies; *synonyms*, containing drug names. Both DrugCentral’s and UMLS’s cross-references to MeSH were used to map DrugCentral internal structure IDs to SemMedDB, ensuring maximum overlap. Disease concepts contained both MeSH and Systematized Nomenclature of Medicine (SNOMED) identifiers that could be mapped to SemMedDB via UMLS cross-references. Some diseases could not be mapped to UMLS, primarily due to the specific nature of the condition, and were discarded. Unmappable conditions included ‘Uremic Bleeding Tendency’, ‘Tonic-Clonic Epilepsy Treatment Adjunct’, and ‘Prevention of Stress Ulcer.’ Highly related diseases were merged to produce a more general disease concept for each treated disease. For example, ‘Vasomotor rhinitis,’ ‘Allergic rhinitis’, ‘Perennial allergic rhinitis’, and ‘Seasonal allergic rhinitis,’ were merged to the single concept ‘Allergic rhinitis.’ For time-resolved analysis, the first approval year for a drug in an indication, provided by DrugCentral, was taken as a proxy for the date of the indication.

### Repurposing Algorithm

A customized version of the PathPredict algorithm [16] utilized in the Repehtio repurposing project [11] was adapted for producing repurposing predictions on the SemMedDB hetnet. This algorithm utilizes Degree Weighted Path Counts (DWPC) as the primary feature for machine learning [17]. These features are based on the various metapaths that connect the source and target node types (in this case Chemicals & Drugs, and Disorders). To aid in the speed of feature extraction, we built a framework (https://github.com/mmayers12/hetnet_ml) based on multiplication of Degree-Weighted adjacency matrices to extract path-counts quickly. The extracted features were then scaled and standardized according to the Rephetio framework. Finally, an ElasticNet regularized logistic regression was performed using the python wrapper (https://github.com/civisanalytics/python-glmnet) for the Fortran library used in the R package glmnet [18]. Hyperparameters were tuned via grid search and once chosen left constant throughout all future runs.

To evaluate the model, the DrugCentral gold standard was partitioned by indication into 5 equal partitions. One-fifth of the indications were withheld during training, and negative training examples were sampled at a rate of ten times the number of positives from the set of non-positive drug-disease pairs. The corresponding TREATS edges for holdout indications were removed from the hetnet before feature extraction in an attempt to limit the model’s ability to learn directly from those edges. The five-fold cross-validations were performed a total of ten times, each with a different random partitioning.

### Time-restricted learning models

The models for the time-resolved networks were trained using the positive gold-standard indications where drug was approved in the years prior to and including the year of the network. Training negatives were selected randomly from the pool of non-positive drug-disease pairs at a rate of ten times the number of positives. After training, the models were then tested on positive indications dated after the year of the network, as well as a proportional number of negatives.

To combine the results of all of the models across the varying network years, the prediction probability for each model was first converted to z-score. This allowed for a cross model comparison of the results. The standardized probabilities for gold-standard drug-disease indications were then grouped according to the difference in years between the network the probability was derived from and the approval year of the drug in the indication. This grouping allowed for the generation of performance metrics for a relative drug approval year. Negative examples were chosen at random from the non-positive set of drug-disease pairs, across all models, at a rate of ten times that of the positives. Area under the receiver operator characteristic (AUROC) and precision recall curves (AUPRC) were then calculated for each of the different time differences from negative 20 to positive 20 years.

### Feature performance analyses

To test the relative importance of each edge type to the model, one of the better performing networks on future indications, 1985, was chosen as a baseline. We performed a ‘dropout’ analysis in which edge instances were removed randomly from the network at rates of 25%, 50%, 75%, and 100% before running the machine learning pipeline. For dropout rates of 25%, 50%, and 75%, the 5 replicates were run with different random seeds, to account for the differences that specific edges may produce when selected for dropout. Performance metrics AUROC and AUPRC of these different dropout results were then compared to the baseline 1985 network model result.

For the edge replacement analysis, the 1985 network was taken as a baseline. Edge instances of a given type were, type by type, replaced with those from the networks of other years starting with 1950 and continuing to present. This produced 15 models for each of the 30 edge types, one for each network year per edge type. For example, for the TREATS edge, all values from the 1985 network were removed and replaced with TREATS edges from the 1950 network and predictions were made, then the TREATS edges were replaced with those from the 1955 network, and so-forth. AUROC and AUPRC results from these modified networks were compared to that of the base 1985 network.

## Results

### 5-fold cross-validation on text-mined data

A hetnet comprised of biomedical knowledge was built from SemMedDB, a database containing subject, predicate, object triples that were text-mined from PubMed abstracts. After data processing steps (see methods) the final network contained 78,400 unique concepts (graph nodes) and 2,470,050 relations (edges) connecting those concepts. These concepts were classified into 6 different types derived from UMLS semantic groups – ‘Chemicals & Drugs’, ‘Disorders’, ‘Genes & Molecular Sequences’, ‘Anatomy’, ‘Physiology’, and ‘Phenomena’. The relationships between the nodes were also classified as one of 30 different edge types, comprised of both a semantic relation and the source and target node types. For example, the relation ‘AFFECTS’ between nodes of type ‘Chemicals & Drugs’ and ‘Anatomy’ is distinct from the relationship ‘AFFECTS’ between nodes of type ‘Chemicals & Drugs’ and ‘Physiology’. In labeling these relations, the node abbreviations are appended to the semantic relation to explicitly differentiate the edge types, e.g. the above examples the labels are ‘AFFECTS_CDafA’ and ‘AFFECTS_CDafPH’ respectively (Table 1, and Supplemental Figure S1, Additional File 1). To train a learning model for compound repurposing, a gold standard of high quality and reliability containing drug-disease indications is required. We used DrugCentral as the source for our gold standard. This open drug database contains a relatively complete, curated list of known indications, with a total of 10,938 unique drug-disease pairs. In mapping these drug and disease concepts to those found in SemMedDB, 3,885 indications were lost due an inability to map the disease condition to a unique concept ID (see methods for examples), and further reductions came due to the merging of highly related disease concepts, resulting in 5,337 unique indications that could be used as true-positives for training and testing purposes.

**Table 1:**
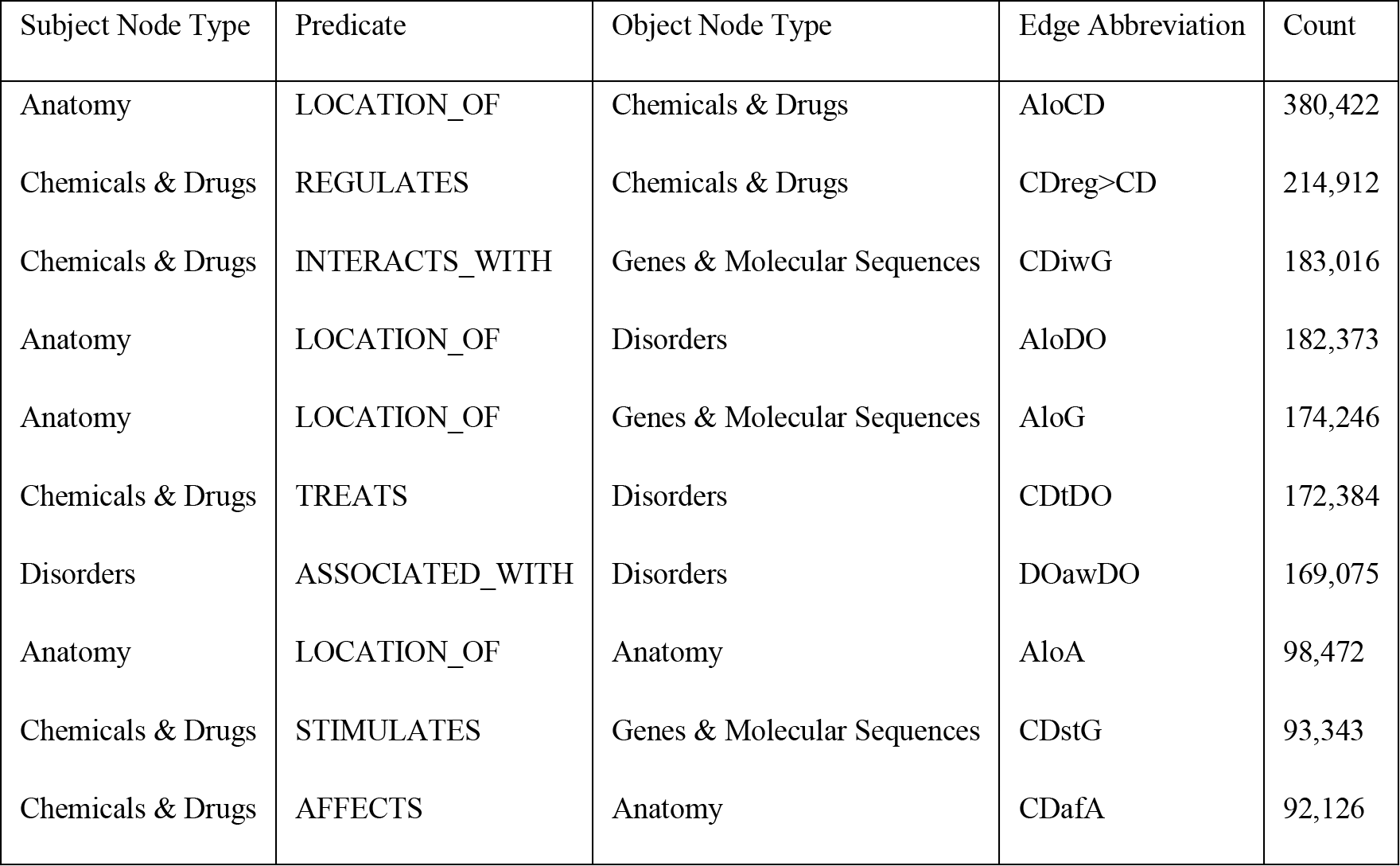
Top 10 Edge Types by Instance Number

After preparation of the hetnet and the gold standard, the utility of this text-mined knowledge base for the prediction of novel drug-disease indications was examined using a modified version of the PathPredict algorithm, utilized by Himmelstein et. al. in the Rephetio drug repurposing project [11]. This paradigm utilizes the degree weighted path count (DWPC) metric, derived from the metapaths that connect different concepts within a network, as the primary features for training the classifier [17]. The remaining features, while comparatively small, are derived from the simple degree values of each edge type for the drug node and the disease node in given drug-disease pair. A 5-fold cross validation was repeated 10 times, each with a random split of the gold standard into training and test sets. The results of the 5-fold cross validation showed excellent results, with an average area under the receiver operator characteristic (AUROC) of 0.95 and average precision (AUPRC) of 0.74 (Figure 1A and 1B). These results are consistent with a very accurate classifier, and comparable to results seen in similar computational repositioning studies [6, 9, 11]. To further evaluate the accuracy of these predictions, the prediction rankings of test set indications were examined for given drugs and diseases (Figure 1C and 1D). The median value for the rank of a positive disease, given a test-set positive drug was 18 out of 740 total diseases. Similarly, when examining the test-set positive diseases, the median rank for a positive drug was 32 out of a possible 1330 examined compounds.

**Figure 1:**
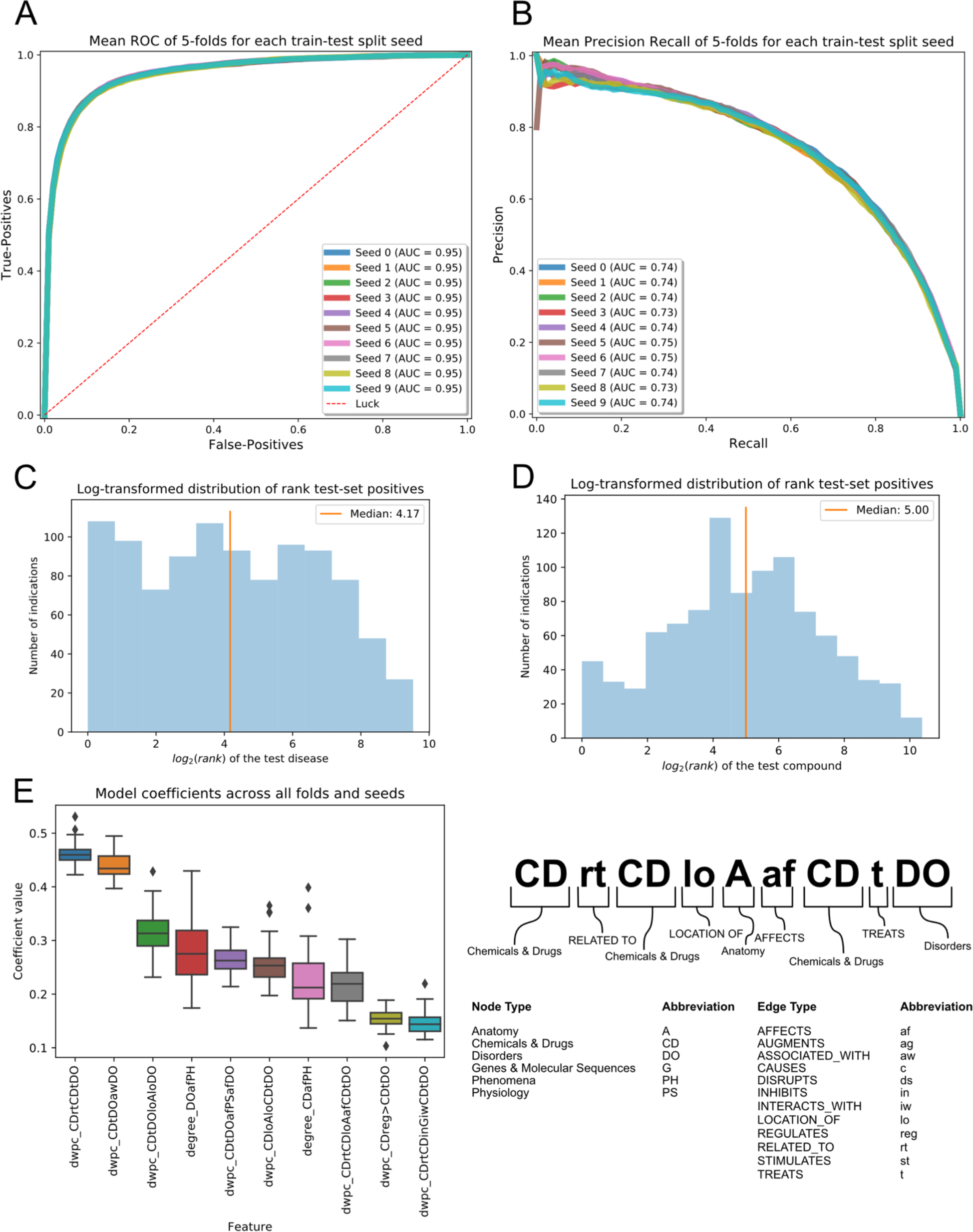
5-fold cross validation results for SemMedDB network using DrugCentral gold standard. **A)** Receiver-Operator Characteristic curve displaying the mean result across 5-folds. Ten different seed values for randomly splitting indications in 5 are compared showing very little variation. **B)** Precision-Recall curve for the mean result across 5-folds, with ten different split seeds displayed. **C)** Histogram of log_2_ transformed rank of true positive disease for a given test-set positive drug, taken from a representative fold and seed of the cross-validation. If a drug treats multiple diseases, the ranks of all diseases treated in the test-set indications are shown. **D)** Histogram of log_2_ transformed rank of true positive drug for a given test-set disease, chosen from same fold and seed as C. If a disease is treated by multiple drugs in the test-set indications, all ranks are included. **E)** (left) Boxplot of 10 largest model coefficients in selected features across all folds and seeds. (right) Breakdown of metapath abbreviations. Node abbreviations appear in capital letters while edge abbreviations appear lower case.

The ElasticNet logistic regression in this analysis used feature selection to reduce the risk of overfitting with a highly complex model. In comparing the models, there was a fairly consistent selection of short metapaths with only two edges that include important drug-drug or disease-disease similarity measures (Figure 1E). These include two related drugs, one of which treats a disease (dwpc_CDrtCDtDO), or two associated diseases, one of which has a known drug treatment (dwpc_CDtDOawDO). However, other metapaths of length 3 which encapsulated drug-drug or disease-disease similarities were also highly ranked. This includes two drugs that co-localize to a given anatomical structure (dwpc_CDloAloCDtDO), two diseases that present in the same anatomical structure (dwpc_CDtDOloAloDO), or diseases that affect similar phenomena (dwpc_CDtDOafPHafDO). In this case anatomical structures could include body regions, organs, cell types or components, or tissues, while phenomena include biological functions, processes, or environmental effects. It is important to again note that these ‘similarity measures’ are purely derived from text-mined relations.

While these results indicate a fairly accurate classifier in this synthetic setting, the paradigm under which they are trained and tested is not necessarily optimal for finding novel drug-disease indications. A cross-validation framework essentially optimizes finding a subset of indication data that has been *randomly* removed from a training set. However, prediction accuracy on randomly removed indications does not necessarily extrapolate to prospective prediction of new drug repurposing candidates. Framing the evaluation framework instead as one of future prediction based on past examples may be more informative. For example, the question ‘given today’s state of biomedical knowledge, can future indications be predicted?’ may more closely reflect the problem being addressed in drug repositioning. The best way to address this question would be to perform the predictions in a time-resolved fashion, training on contemporary data and then evaluating the model’s performance on an indication set from the future.

### Building time-resolved networks

To facilitate a time-resolved analysis, both the knowledge base data and the training data need to be mapped to a particular time point. Each triple in SemMedDB is annotated with a PMID, indicating source abstract of this text-mined data. Using the PMID, each triple, corresponding to an edge in the final network, can be mapped to a specific date of publication. The DrugCentral database also includes approval dates from several international medical agencies for the majority of the drugs. By filtering the edges in the network by date, an approximate map of the biomedical knowledge of a given time period can be produced. Therefore, we generated multiple networks, each representing distinct time-points. We then applied the machine learning pipeline to each of these networks to evaluate the expected performance on future drug-disease indications. Combining these sources of time-points for the network serves to replicate the paradigm of training a machine learning model on the current state of biomedical knowledge, evaluating its ability to predict what indications are likely to be found useful in the future.

Knowledge networks were built in a time-resolved fashion for each year, starting with 1950 and continuing until the present. This was accomplished by removing edges with their earliest supporting PMID dated after the desired year of the network. If either a drug or a disease from a known gold standard indication was no longer connected to any other concept in the network, the indication was also removed from the training and testing set for that network year. Examining the trends of the networks constructed for the various timepoints, the number of nodes and edges always increased, but edges increased more quickly with later timepoints producing a more connected network than earlier (Figures 2A and 2B).

**Figure 2:**
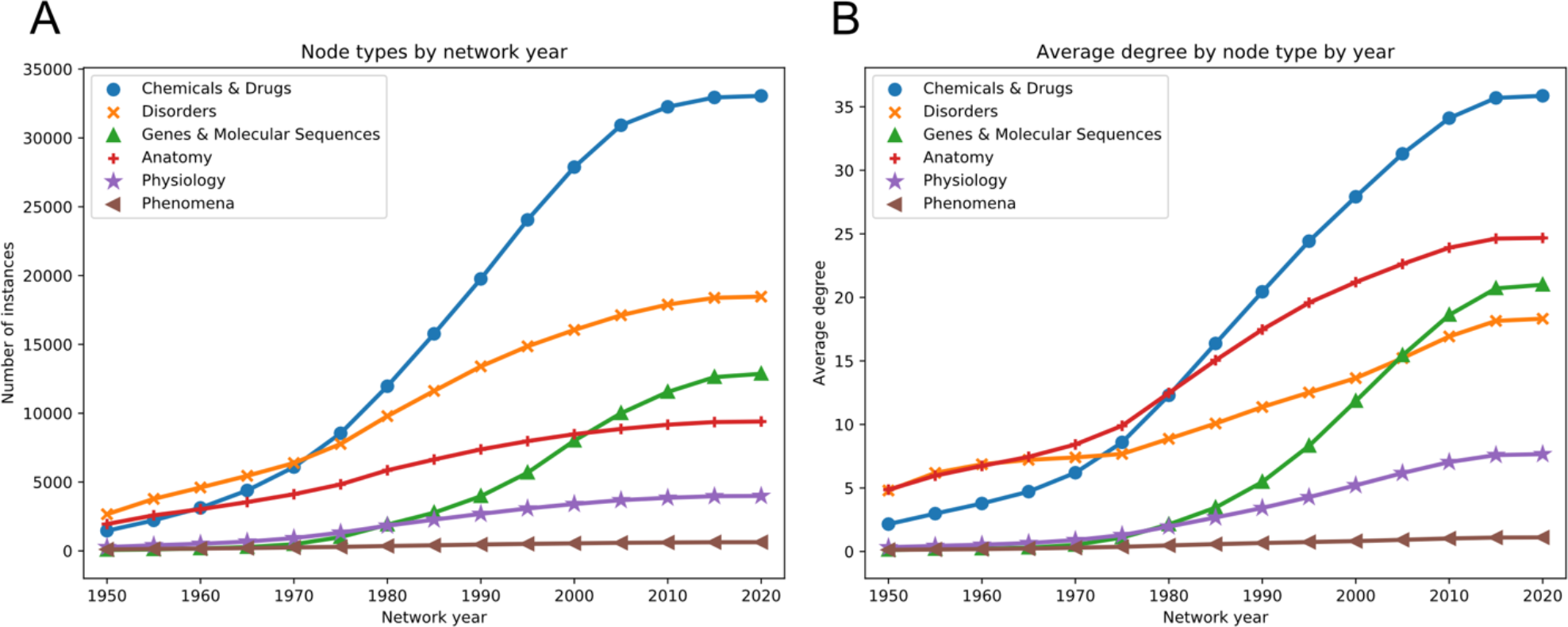
Time-resolved network build results. **A)** Number of nodes of a given type by network year. **B)** Average node degree for each node type across all network years.

The number of indications that could be mapped to a given network year increased quickly at first but rose much more slowly in the later years of the network, even though the total number of concepts in the network continued to increase. For the majority of the years of the network, the split between current and future indications remained at a ratio of around 80% current and 20%, ideal for a training and testing split. However, after the year 2000, the number of mappable future indications continued to diminish year after year, reducing the test set size for these years (Supplemental Figure S2, Additional File 1).

### Machine learning results

The performance of each model against a test set of future indications steadily increased from the earliest time-point until the 1987 network. The AUROC metric saw continual increases over the entirety of the network years, though these increases occurred more slowly after the 1987 network (Figure 3A). Looking at average precision, this metric peaked at the 1987 timepoint with a value of 0.492, and then fell sharply at 2000 and beyond, likely due to the diminished number of test-set positives. The AUROC of this peak average precision time point of 1985 was 0.822. These peak performance metrics fall far below those found via 5-fold cross-validation indicating an inherent limitation in evaluating models via this paradigm.

**Figure 3:**
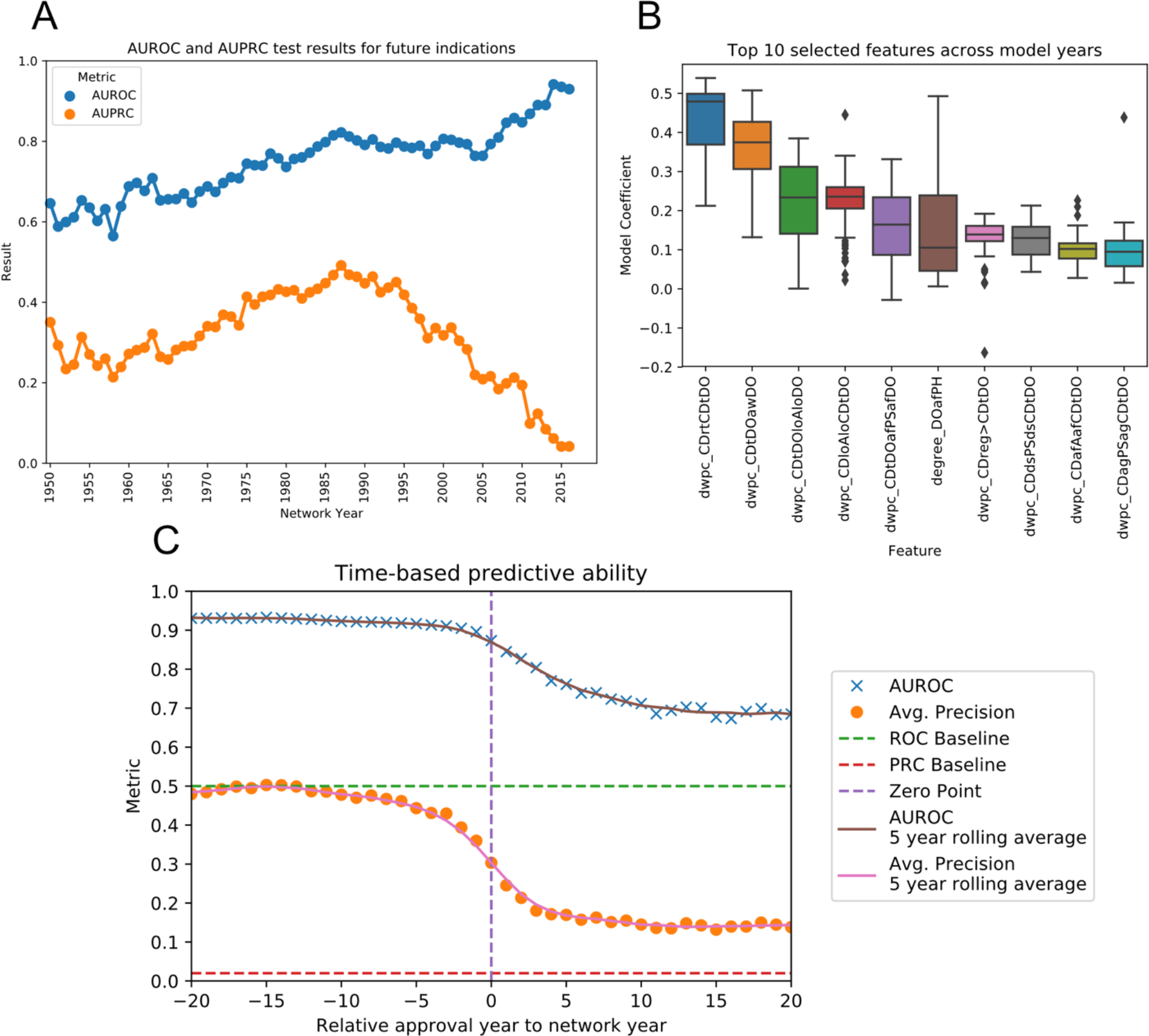
Machine learning results for the time-resolved networks. **A)** Performance metrics for the test-set (future) indications across the different network years. Only drugs approved after the year of the network are included in the test-set, while those approved prior are used for training. **B)** Box plots of the values of the model coefficients across all of the different network years. The top-10 coefficients with largest mean value across all models are shown. **C)** AUROC and AUPRC data for indications based on their probabilities, split by the number of years between drug approval date and the year of the network. Values to the left of the Zero Point are indications approved before the network year thus part of the training-set, while those to the right are part of the test-set. Probabilities for all drug-disease pairs were standardized before combining across models. Points are given for each data point, while lines represent a 5-year rolling average of metrics.

Similar to the cross-validation results, the models favored metapaths that represented drug-drug and disease-disease similarity (Figure 3B). Specifically, the metapaths of type ‘Chemical & Drug - TREATS - Disorder - ASSOCIATED WITH - Disorder’ (dwpc_CDtDOawDO) and ‘Chemical & Drug - RELATED_TO - Chemical & Drug - TREATS - Disorder’ (dwpc_CDrtCDtDO) had the highest weights across almost all models. One difference found from the cross-validation results is the appearance of the ‘Physiology’ metanode in two of the top selected metapaths, one connecting two diseases through common physiology, and one connecting two drugs that both augment a particular physiology. Model complexity was also diminished compared to those seen in during cross-validation, with the majority of models selecting less than 400 features, or 20% of the total available (Supplemental Figure S3, Additional File 1).

Finally, one question to explore is whether or not there is a temporal dependence on the ability to predict indications. For example, is there better performance on drugs approved 5 years into the future rather than 20, since one only 5 years pre-approval may already be in the pipeline with some important associations already known in the literature. To answer this, the results from all network years were combined via z-scores. Grouping indications by approval relative to the year of the network allowed for an AUROC metric to be determined for different timepoints into the future (Figure 3C). This analysis revealed that there is still a substantial predictive ability for drugs approved up to about 5 years into the future. However, after 5 years, this value quickly drops to a baseline of .70 for the AUROC and .15 for the average precision. These results indicate a temporal dependence on the ability to predict future indications, with the model being fairly inaccurate when looking far into the future.

### Edge dropout confirms importance of drug disease links

Many other efforts in computational repositioning have found that emphasis on drug-drug and disease-disease similarity metrics results in accurate predictors [6, 19, 20]. To further investigate the types of information most impactful in improving the final model, an edge dropout analysis was run. The 1985 network was chosen as a base network for this analysis both due to its relatively strong performance on future indications and its centralized time point among all the available networks. By taking each edge type, randomly dropping out edge instances at rates of 25%, 50%, 75% and 100%, and comparing the resulting models, the relative importance of each edge type within the model could be determined. The edge that was found to have the largest impact on the resulting model was the ‘Chemicals & Drugs - TREATS - Disorders’ edge, reducing the AUROC by .098 (Figure 4A). This result reinforces the idea that drug-disease links, particularly those with a positive treatment association, are highly predictive in repositioning studies. The drug-drug (‘Chemicals & Drugs - RELATED_TO - Chemicals & Drugs’) and disease-disease (‘Disorders - ASSOCIATED_WITH - Disorders’) similarity edges were the next two most impactful edges on the overall model, both showing decreases of .015 in the AUROC when completely removed. Overall, however most edges showed very little reduction in AUROC, even at 100% dropout rate. This could indicate a redundancy in important connections between drugs and diseases that the model can continue to learn on even when partially removed.

**Figure 4:**
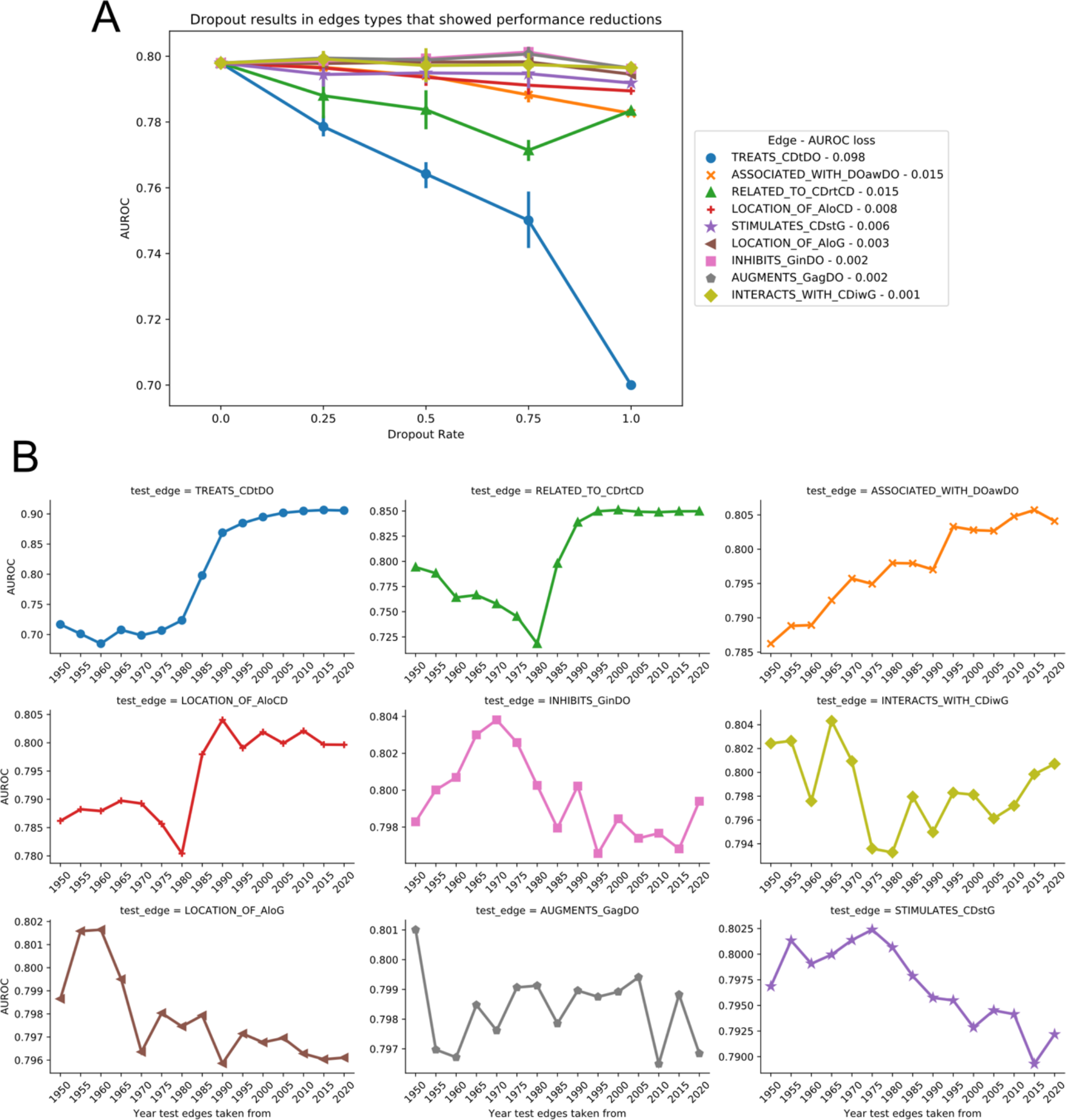
Analysis of edge type importance to the overall model. **A)** Edge dropout analysis showing the reduction in AUROC metric when the edges are dropped out at rates of 25, 50, 75, and 100%. Error bars indicate 95% confidence interval over 5 replicates with different seeds for dropout. The 9 edge types that had the greatest reduction from 0 to 100% dropout are displayed. **B)** Edge replacement analysis showing changes in AUROC when edges are replaced with those of the same type from another year’s network. The top 9 edges that showed greatest loss in performance in the dropout analysis between 0 and 100% dropout are displayed.

### Time-resolved edge substitution confirms edge importance

While dropout identifies the most important associations between concepts to this predictive model, this does not necessarily confirm that more data of these types will improve the model’s results. To simulate this the impact of the assimilation of new knowledge of a specific type, an edge replacement analysis was performed on the 1985 network. This process allowed for the examination of how accumulating new real-world data of a given type might affect the model. By taking a specific edge type and replacing all the edges of that type with those from the other network years from 1950 to 2015, the potential effect of gathering more data of these specific types over time could be examined. Similar to the dropout analysis, the target edge of ‘Chemicals & Drugs - TREATS - Disorders’ had the greatest effect on the model’s performance, showing an increase of .108 when replaced with the most current version of the edge (Figure 4B). Similarly, the AUROC showed a large loss of .081 when replaced with values from 1950. The drug-drug and disease-disease similarity edges also showed significant performance increases when replaced with contemporary values, while decreasing performance in performance when replaced with 1950 values. While the three edges that produced the greatest decrease in performance during the dropout analysis also had the biggest benefit when adding future edges, not all behaved in this manner. For example, the edge ‘Anatomy - LOCATION_OF - Chemicals & Drugs’ showed the fourth largest decreases in performance during edge dropout analysis. When using past versions of this edge type with the 1985 network, the performance did have a measurable decrease in AUROC of .012, however current versions of this edge type only improved the score by .002. Conversely, the edge ‘Physiology - AFFECTS - Disorders’ showed little to no performance loss during the dropout analysis and indeed showed little performance change when using past versions of the edge (Supplemental Figure S4, Additional File 1). However, this edge showed substantial increase of .012 AUROC when using contemporary versions of the edge. Finally, some edge types like ‘Genes & Molecular Sequences - ASSOCIATED WITH - Disorders’ actually performed slightly better with past version or future versions of the edge, when compared 1985 version of the edge, with an increase in AUROC of .004 with contemporary edges and an increase of .011 with edges from 1950 (Supplemental Figure S5, Additional File 1). This further underscores the idea that a time-resolved analysis provides a more complete picture of the important components to a learning model.

## Discussion and Conclusions

While a text-mined data source, SemMedDB performed very well when using the metapath-based repositioning algorithm from Rephetio and trained and tested against a DrugCentral derived gold standard. However, performing well in a cross-validation does not necessarily lead to a large number of real-world repositioning candidates. This evaluation paradigm essentially trains the learning model to identify indications that are currently known but simply withheld from a dataset. In the real world, the problem solved by computational repositioning is more closely aligned to attempting to predict new indications that are not already known at this current time-point. Our use of time-resolved knowledge networks has allowed us to replicate this paradigm and expose a marked reduction in performance when a model is tested in this fashion. Time separation is a long-used practice to combat overfitting in data mining [21] and our application of this practice to compound repositioning may help explain some of the discrepancy between model performance and the number of repositioning candidates successfully produced through computational repositioning.

We believe that this method for evaluating a repositioning algorithm in a time-resolved fashion may more accurately reflect its ability to find true repurposing candidates. Identifying algorithms that perform well at predicting future indications on the time-resolved networks presented in this paper may yield better results when translating retrospective computational analyses to the prospective hypothesis generation. As these networks are built around text-mined data, predictive performance may be enhanced by utilizing high-confidence, curated, data sources for computational repositioning. The original date of discovery for a given data point has shown itself to be an important piece of metadata in evaluating a predictive model. Ensuring curated data sources are supported by evidence that can be mapped back to an initial date of discovery functions to enhance the utility of the data in predictive models such as these. Finally, this temporal analysis again supports the notion that drug and disease similarity measures as well as direct associations between these concepts are still the most important pieces of data in generating a predictive model. Further enhancing our understanding of mechanistic relationships that these concepts will likely result in further increases to computational repositioning performance.

## List of abbreviations

Hetnet: heterogeneous network
NLP: Natural Language Processing
SemMedDB: Semantic Medline Database
PMID: PubMed Identifier
DWPC: Degree Weighted Path Count
AUROC: Aera Under the Reciever Operator Curve
AUPRC: Area Under the Precision Recall Curve (aka average precision)
UMLS: Unified Medical Language System
MeSH: Medical Subject Headings

## Ethics approval and consent to participate

Not applicable.

## Consent for publication

Not applicable.

## Availability of data and materials

Data for SemMedDB hetnet building: The SemMedDB database used to build the heterogeneous network analyzed in this study are is available here: https://skr3.nlm.nih.gov/SemMedDB/index.html

The UMLS Metathesaurus used for identifier cross-referencing are available https://www.nlm.nih.gov/research/umls/licensedcontent/umlsknowledgesources.html

These data are provided by the UMLS Terminology Service, but restrictions apply to the availability of this data, which were used under the UMLS Metathesaurus License. https://www.nlm.nih.gov/databases/umls.html#license_request ^[14]^

Data for gold standard: The DrugCentral database used to build the gold standard for this study is freely available from DrugCentral under the CC-BY-SA-4.0 license. http://drugcentral.org/ ^[15]^

Source code to download the above datasets and reproduce the analysis found in this current study is available on GitHub in the following repository. https://github.com/mmayers12/semmed

## Author’s contributions

MM developed the network building pipeline, adapted the machine learning algorithm for use with SemMedDB, and wrote the majority of the manuscript. TL developed the DrugCentral gold standard and designed the feature performance analysis experiments. NQ organized graph data and participated in design of the experiments. The research was performed under the advice and supervision of AS. All authors read and approved the final manuscript.

## Acknowledgements

N/A

## Additional files

Additional File 1: Supplemental Figures

