## Additional File 1 for "Time-resolved compound repositioning predictions on a text-mined knowledge network"

### Supplemental Figures

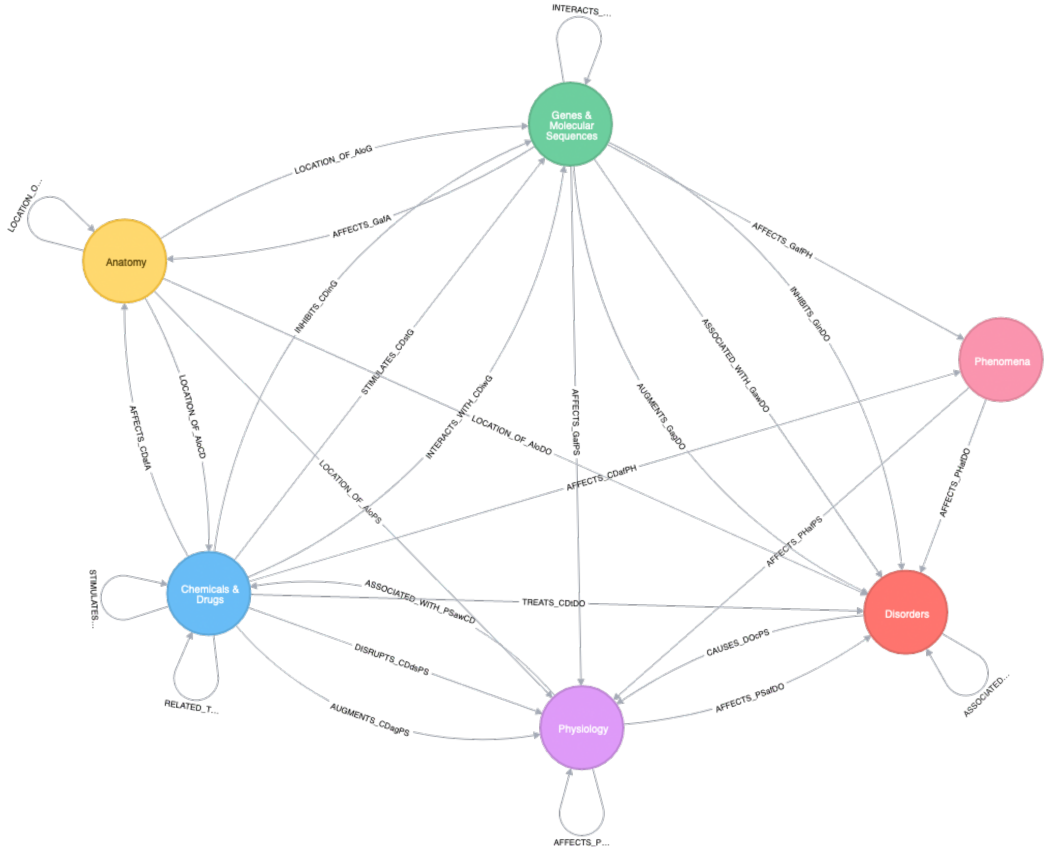


**Supplemental Figure S1)** Metagraph of SemMedDB hetnet data model.

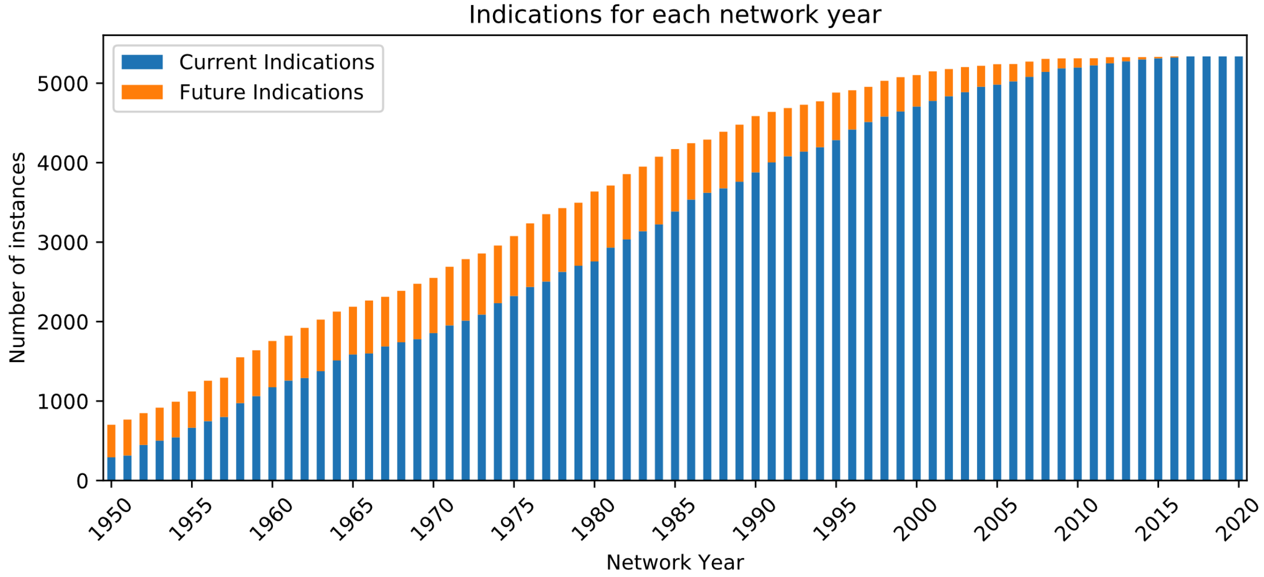


**Supplemental Figure S2)** Number of indications mappable to each network year, split by approval year, with current indications mean those approved up-to and including the network year, and future those approved after the network year.

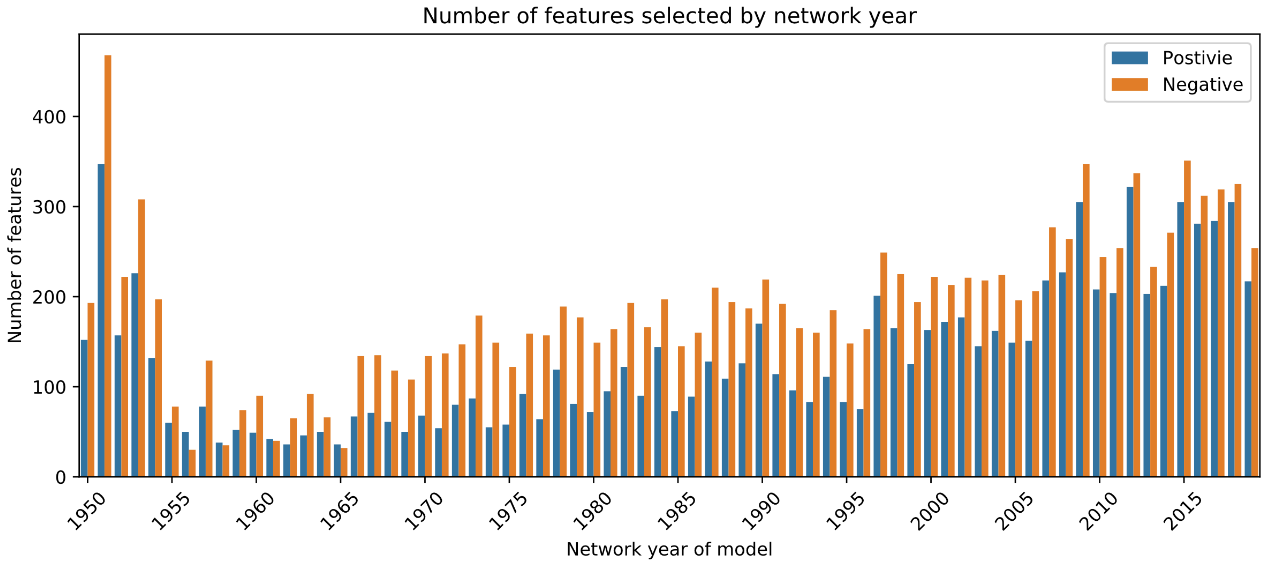


**Supplemental Figure S3)** Number of features selected in the models for each of the different network years.

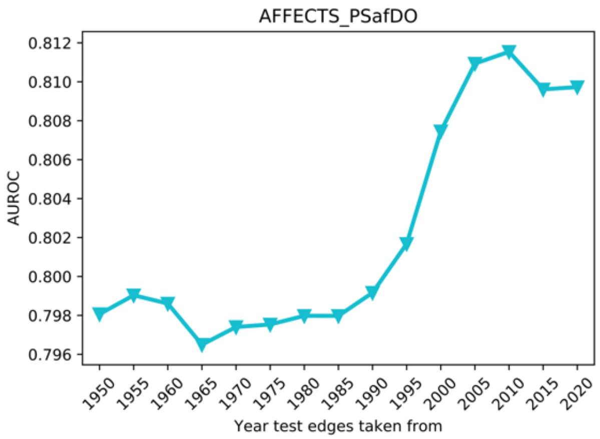


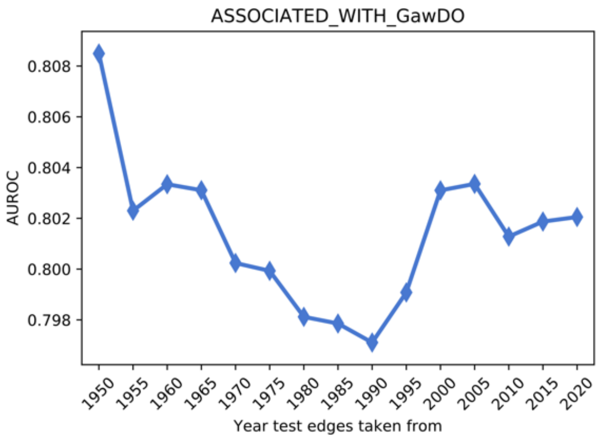


**Supplemental Figures S4 & S5)** Edge substitution analysis results for edges AFFECTS_PSafDO and ASSOCIATED_WITH_GawDO respectively.
